# Adrenergic Reprogramming of Preexisting Adipogenic Trajectories Steer Naïve Mural Cells Toward Beige Differentiation

**DOI:** 10.1101/2023.08.26.554950

**Authors:** Kathleen Desevin, Briana N. Cortez, Jean Z Lin, Dechen Lama, Matthew D. Layne, Stephen R. Farmer, Nabil Rabhi

**Affiliations:** Department of Biochemistry and Cell biology, Boston University Chobanian & Avedisian School of Medicine, Boston, MA, USA

**Keywords:** Adipose tissue, Beige Adipogenesis, Thermogenesis, Progenitor Cells, Mural Cells, Adrenergic, Lineage Tracing, Obesity, Metabolic Diseases, Cell-Cell Communication

## Abstract

In adult white adipose tissue, cold or β3-adrenoceptor activation promotes the appearance of thermogenic beige adipocytes. Our comprehensive single-cell analysis revealed that these cells arise through the reprogramming of existing adipogenic trajectories, rather than from a single precursor. These trajectories predominantly arise from SM22-expressing vascular mural progenitor cells. Central in this transition is the activation of Adrb3 in mature adipocytes, leading to subsequent upregulation of Adrb1 in primed progenitors. Under thermoneutral conditions, synergistic activation of both Adrb3 and Adrb1 recapitulates the pattern of cold-induced SM22+ cell recruitment. Lipolysis-derived eicosanoids, specifically docosahexaenoic acid (DHA) and arachidonic acid (AA) prime these processes and in vitro, were sufficient to recapitulate progenitor cells priming. Collectively, our findings provide a robust model for cold-induced beige adipogenesis, emphasizing a profound relationship between mature adipocytes and mural cells during cold acclimation, and revealing the metabolic potential of this unique cellular reservoir.

## INTRODUCTION

Adipose tissue (AT) is a highly complex organ comprising mature adipocytes, immune cells, and stromal vascular cells. AT possesses a remarkable ability to adapt to environmental and dietary cues by dynamically altering its morphology and metabolic capacity. Under specific stimuli, such as cold exposure or β-3-adrenergic (Adrb3) activation, white adipose tissue (WAT) can undergo a transformation to acquire a thermogenic phenotype characterized by the appearance of mature adipocytes expressing uncoupling protein 1 (UCP1). These cells, often referred to as beige or brite adipocytes, exhibit energy-burning capabilities similar to those of brown adipose tissue (BAT) but possess a distinct molecular and developmental trajectory from BAT^1,2,3^. Enhancing beige adipocyte recruitment and activation, while suppressing white adipocytes, holds great potential for combating obesity and its associated metabolic complications. Indeed, activation of thermogenesis promotes a shift in energy expenditure among obese and type 2 diabetes (T2D) individuals, leading to improved glucose and lipid clearance to support thermogenesis^4,5,6,7^. Additional studies have demonstrated that promoting beige adipocytes prevent high-fat diet-induced insulin resistance, weight gain, WAT inflammation, fibrosis, and hepatic steatosis, resulting in enhanced metabolic indicators such as insulin sensitivity and euglycemia^7,8,9,10,11,12^. In human subjects, ADRB3 activation and cold acclimation stimulate thermogenic adipocytes, ameliorating obesity’s detrimental outcomes, including T2D, dyslipidemia, and cardiometabolic diseases^7,13,14,15,16^. Furthermore, adult human BAT from the supraclavicular region harbors adipocytes displaying a molecular signature akin to murine beige adipocytes^16,17,18,19^.

In adult mice, cold exposure triggers the emergence of beige adipocytes through two distinct mechanisms: de novo beige adipogenesis from adipose progenitor cells (APCs) and the activation of “dormant beige adipocytes” ^20^. Notably, APCs expressing surface markers Pdgfrα and Sca1 have been demonstrated to differentiate into both white and beige adipocytes^21,22^. Within the Pdgfrα+; Sca1+ progenitor cells, Cd81 expression has been associated with APCs beiging potential^23^. Lineage tracing studies have revealed that progenitors expressing Acta2, Pax3, Pdgfrβ, or Myh11 in the stromal vascular fraction (SVF) of WAT are the source of beige adipocytes following cold exposure over timeframes ranging from 7 to 21 days^20,21,24,25^. A recent model proposes that all APCs in subcutaneous WAT belong to the Pdgfraα+ lineage, giving rise to two distinct types of committed preadipocytes: Pdgfrβ− preadipocytes, capable of differentiating into beige adipocytes, and Pdgfrβ+ preadipocytes, which can develop into unilocular white adipocytes^26,27^. Characterization of WAT from Pdgfrα reporter mice identified two distinct cell populations: Lin−:CD29+:CD34+: Sca-1+:CD24+ (CD24+) and Lin−:CD29+:CD34+: Sca-1+:CD24− (CD24−), which serve as adipocyte precursors. The CD24+ cells were found to give rise to the CD24− population in vivo, suggesting that CD24+ adipocyte progenitors progress toward the adipocyte lineage as CD24 expression is lost, generating CD24− preadipocytes^21^. Intriguingly, clonal examination of cultured adipose-derived stromal cell lines suggests the presence of committed beige precursors within adult WAT, separate from white adipocyte precursors^28,17,12^. However, it remains unclear whether adipose tissue hosts functionally distinct beige and white adipocyte precursors or unique bipotent precursors whose commitment to the beige adipocyte lineage is influenced by cold-associated signals.

In this study, we leveraged single-cell sequencing to uncover that identical progenitor cell populations exist within the adipose tissue depot, regardless of the beiging agent used. We discovered that most of the progenitor cells emerge from a single mural cell type expressing transgelin (SM22). Moreover, our investigation revealed that upon cold exposure, lipolysis through β3-AR activation promotes the beige reprogramming or priming of SM22+ progenitor cells, leading to their cellular expansion. These naïve cells may be considered to be in an “undetermined” state. However, as they progress toward the beige lineage, SM22+ cells begin to express β1-adrenergic (β1-AR) receptors. Activation of β1-AR is essential for the recruitment and subsequent differentiation of these cells into beige adipocytes.

## RESULTS

To delineate early cellular adaptations during the initial stages of beige adipogenesis, we submitted to single-cell sequencing (scRNA-seq) lineage-negative (Lin−) stromal cells isolated from inguinal WAT of adult mice treated with vehicle, Adrb3 agonist (CL 316,243; CL−) or cold for 3 days. Unsupervised clustering resulted in the identification of 16 unique clusters (**Fig. 1A**). Subsequent analysis of integrated data revealed a host of cell populations. Among these were three subpopulations of mesenchymal stem cells (MSC) subgroups (MSC1, MSC2a, and MSC2b), where MSC2a and MSC2b displayed significant overlap in their gene markers program. We also identified two subsets of interstitial cells (ICa, ICb), smooth muscle cells (SMC), and two distinct subpopulations of the endothelial cells (EC1 and EC2). An additional subgroup of cells exhibited similarities to non-SMC mural cells annotated here as mural cells. Interestingly, three cell populations demonstrated shared markers of EC and MSC (MSC-EC-L1, 2, 3). Furthermore, we identified neuronal cells, Schwann cells, and a highly proliferative cell type. All MSC, IC, EC, and mural cells exhibited expression of canonical stromal markers such as Ly6a (Sca1) and Cd34. Pdgfrα was restricted to MSC1, MSC2a, and ICs, while Pdgfrβ primarily appeared in MSC1, SMC, mural cells, and mural-like cells (**Fig. S.1A**). Interstitial fibroblast cell markers including Pi16 and Dpp4 were enriched in two discernable subgroups, distinguished by Dmkn and Krtdap expression (**Fig. 1B, Table S1**). Importantly, at the selected single-cell analysis resolution each population expressed cell-specific markers (**Fig. 1B, S.1B, Table S1**). MSC-EC cell groups, despite expressing some unique cell markers, also demonstrated shared EC and MSC marker expression, suggesting their intermediary nature (**Fig. 1D**).

**Figure 1:**
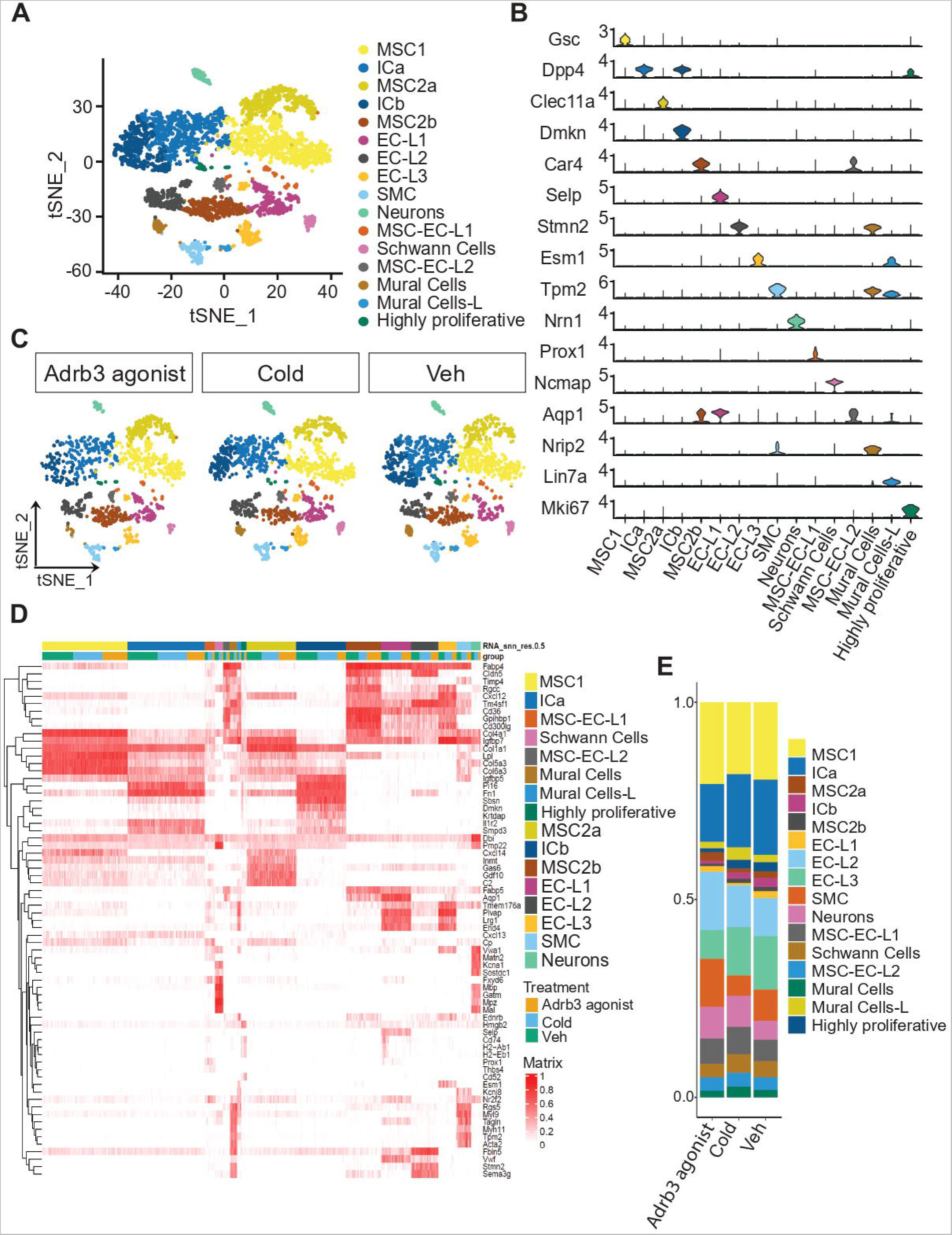
Sustained Cellular Heterogeneity in WAT Unaltered by β3-Adrenergic or Cold Stimulation. (**A)** t-SNE plots showing different Lin-stromal vascular cell types in iWAT isolated from mice treated with Vehicle, CL−316,243 or exposed to 4°C for 3 days. Colors represent 16 different clusters determined by unbiased clustering across the 4879 cells that passed the quality control. MSC: mesenchymal stem cells, IC: interstitial cells, EC-L: endothelial-like cells, SMC: smooth muscle cells. **(B)** Violin plots showing the expression levels and distribution of representative marker genes of each cluster. The y-axis is the log-scale normalized read count. **(C)** Side-by-side t-SNE plots of iWAT Lin-stromal vascular cell types from mice treated with vehicle, CL−316,243 and exposed to 4°C for 3 days. **(D)** Heatmap of the top differentially expressed genes. **(E)** Bar plot showing the percentage of cells per cluster in each treatment.

Unexpectedly, independent analyses of vehicle, CL−, and cold-treated scRNA-seq datasets unveiled identical cell populations (**Fig. 1C, 1D, S.3C**). Further investigation indicated variations in cell numbers per cluster across different treatments (**Fig. 1E**). Interestingly, cell numbers per cluster were significantly changed between cold-exposed and CL− treated mice. The cold-exposed mice displayed a noticeable reduction in the proportion of MSCs compared to CL− treated mice, whereas IC, mural cells, and MSC-EC cells increased under cold exposure. These data imply either that a beige precursor cell population already exists within the tissue before adrenergic stimulation, or that a specific subgroup of cells may be primed to transition towards beige differentiation upon stimulation.

We next pursued an exhaustive cell state analysis. This required the exclusion from the dataset the cells that were predicted to belong to the WAT lineage in our initial analysis including schwann cells and SMC. The resultant pool of cells was then re-clustered at an elevated resolution (Res = 0.6) (**Fig. 2A**), enabling the identification of 17 distinct cell states and types (**Fig. S.2A, S.2B, S.2C**). When the clustering resolution was increased further, the identified cell states/types remained either consistent across conditions or lead to unstable clustering (**Fig. S2D, S2E**). Prior to pseudotime analysis, we performed an imbalance scoring. This allowed us to examine whether the distribution of cells according to the treatment label deviated from the overall cell distribution. Although minor regions of imbalance surfaced between treatments, the main trajectory was preserved across conditions. This suggested that the cellular trajectory was unaffected by alterations introduced by CL− and cold exposure, compared to the vehicle-treated mice (**Fig. 2B**). Subsequently, we undertook a concurrent trajectory analysis for all treatments. This strategy enabled us to uncover the existence of nine potentially distinct lineages within WAT (**Fig. 2C**). While these lineages bore similarities across different conditions, a detailed analysis of the pseudotime distribution along each lineage revealed a significant redistribution of cell types/states. Specifically, substantial alterations in the distribution of cells along lineages were observed when comparing cells isolated from mice subjected to cold exposure or CL− treatment with the vehicle-treated group. This observation was also evident when comparing the total number of cells predicted to contribute to each lineage (**Fig. S2F**).

**Figure 2:**
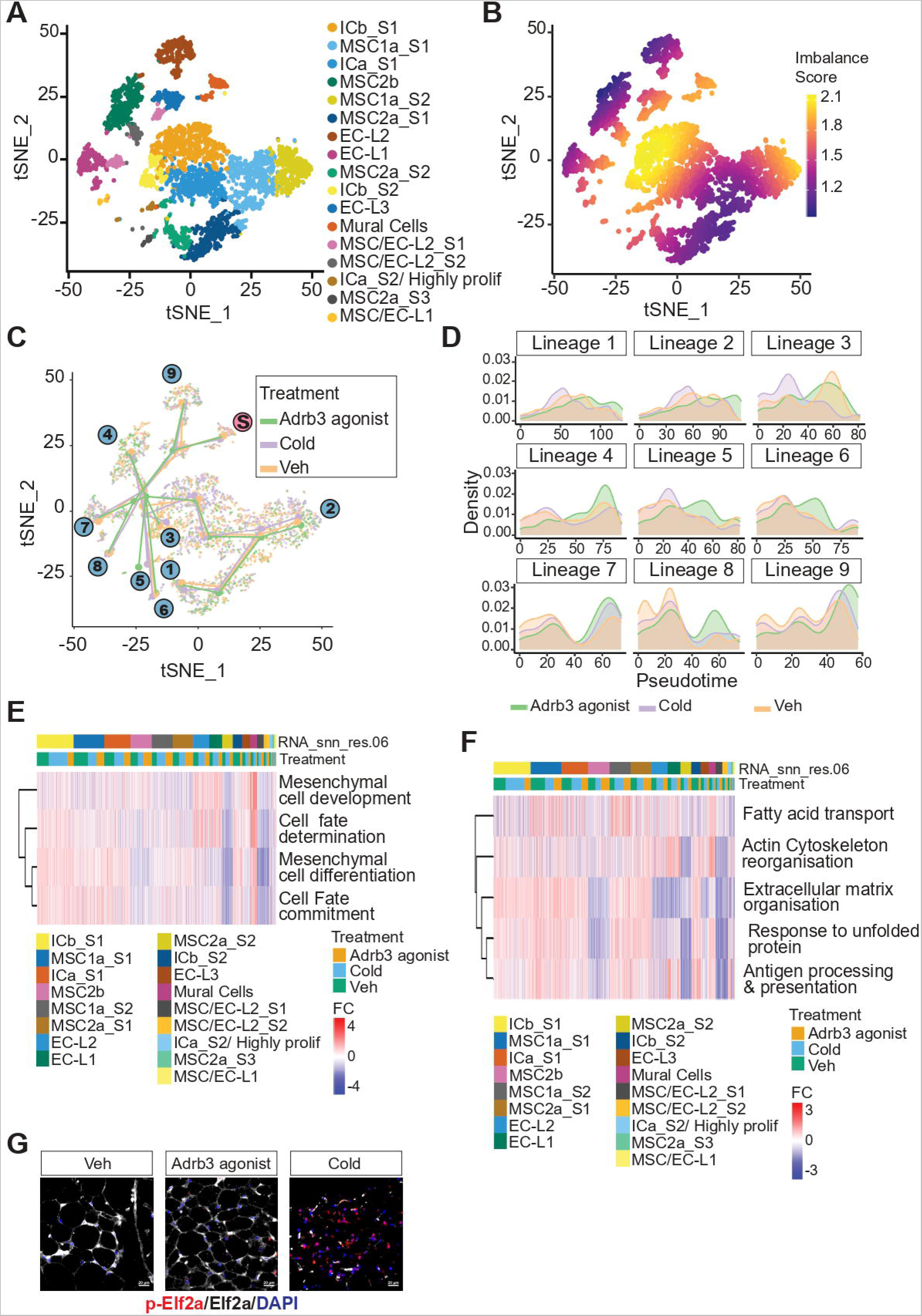
Shift of Cell Trajectory in Response to Cold Exposure and β3-Adrenergic Stimulation. **(A)** t-SNE plots showing unbiased re-clustering of iWAT progenitor cell type states. Colors represent 17 different clusters MSC: mesenchymal stem cells, IC: interstitial cells, EC-L: endothelial-like cells, S; state. **(B)** t-SNE plots showing differential topology in single cell states between mice treated with vehicle, CL−316,243 and exposed to 4°C for 3 days. **(C)** t-SNE plot of inferred trajectories of single cell type states for each treatment. **(D)** Heatmap of enriched pathways (biological process) associated with mesenchymal cell development and fate decision. **(E)** pseudotime distributions per condition along the trajectory of each lineage. **(F)** Heatmap of top gene ontology pathways (biological process) associated with genes differentially expressed in cold exposed samples. **(G)** Representative image of eIf2a and p-elf2a immunostaining scale bars 20 μm.

Intriguingly, we found that all clusters were predicted to emanate from a single subpopulation of mural cells, corroborating previous studies (**Fig 2B**). Further gene ontology analysis revealed an enrichment of pathways tied to cell fate determination, mesenchymal cell development, and cell fate determination specifically linked to this mural cell population (**Fig. 2E**).

Our analysis identified notable differences in cell contributions between the cold-exposed and CL− treated groups. This supported the hypothesis that cold exposure and CL− treatment stimulate distinct dynamics within the APC population. To better understand these differences, we performed a differential gene expression analysis across each lineage while ordering them based on hierarchical clustering under cold conditions (**Fig. S2G**). We found that several gene groups displayed different patterns across conditions and clusters. We subsequently performed a gene set enrichment analysis on these differentially expressed genes between the conditions (**Fig. 2F**). Terms related to fatty acid transport, actin cytoskeleton reorganization, extracellular matrix organization, response to unfolded protein, and antigen processing & presentation were highly enriched across treatments (**Fig. 2F**). Pathway signature scoring demonstrated a cell-type and treatment-dependent enrichment of each pathway (**Fig S2H**). Remarkably, we found that genes associated with extracellular matrix organization and antigen processing & presentation were downregulated in cell types sourced from cold-treated mice. In contrast, genes related to fatty acid transport, actin cytoskeleton reorganization, and response to unfolded protein (UPR) were upregulated. These findings imply that these five pathways chiefly account for the differences observed between conditions. Consistent with our transcriptomic findings, we found a substantial increase of p-eIf2a immunostaining in response to cold compared to CL− or vehicle treatment (**Fig. 2G**).

Collectively, our data support a model where beige adipocytes originate from the reprogramming of the preexisting adipogenic trajectory, rather than the activation of a unique beige adipogenic trajectory. Furthermore, our single-cell data analysis revealed that cold exposure and β3-adrenergic stimulation each promote distinct cellular dynamics.

To investigate whether cells are indeed reprogramed toward the beige fate, we assessed gene expression of thermogenic genes on isolated progenitor cells from mice treated with cold, CL−, or vehicle for 3 days (**Fig. S.3A**). Remarkably, an enhancement in thermogenic markers was elicited within progenitors by both the cold and CL− treatments (**Fig. 3A**). Subsequent in vitro differentiation assays, employing the standard white adipogenic cocktail, unveiled that progenitor cells isolated from either cold- or CL− exposed mice differentiated into beige adipocytes while those isolated from the control group developed into white adipocytes (**Fig. 3B**). This suggests a “priming” or predisposition of progenitor cells isolated from mice treated with CL− or exposed to cold to adopt the beige adipogenic fate.

**Figure 3:**
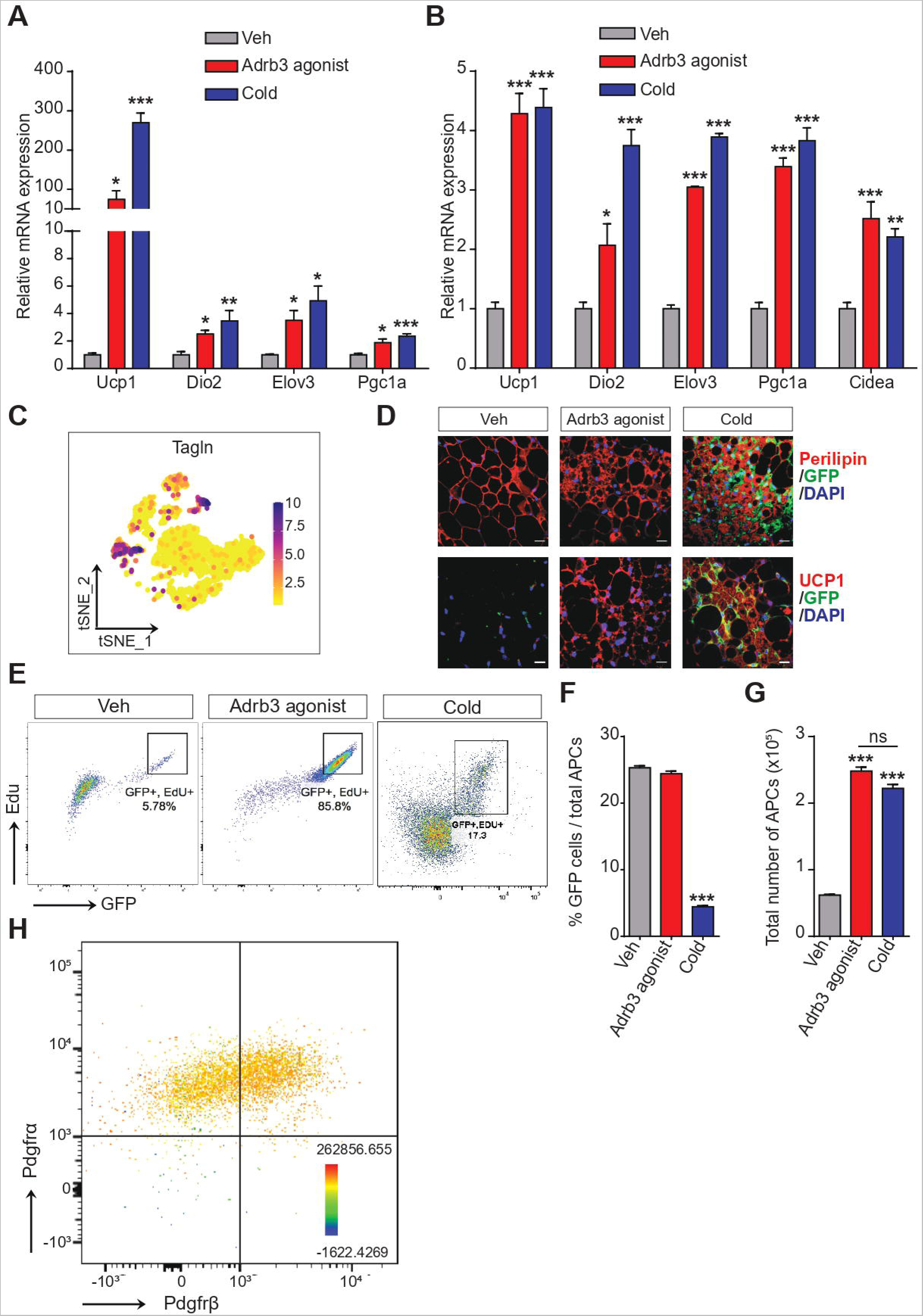
SM22 Expressing Mural Cells are Primed into the Beige adipogenic cells but only cold promote their Recruitment. **(A)** Gene expression of thermogenic markers in isolated Lin-Sca+ cells from mice treated with vehicle, CL−316,243 and exposed to 4°C for 3 days. **(B)** Gene expression of thermogenic markers in vitro differentiated Lin-Sca+ cells from mice treated with vehicle, CL−316,243 and exposed to 4°C for 3 days. **(C)** t-SNE plot of Tgln expression level and distribution. **(D)** Immunostaining in SM22-rtTA-cre;mTmG mice. **(E)** Representative scatter plot of flow cytometry analysis of EdU incorporation in GFP+, Lin-, Cd31-, Sca+ cells. **(F)** Quantification of GFP+ Lin-, Cd31-, Sca+ cells relative to total Lin-, Cd31-, Sca+ cells. **(G)** Absolute counts of Lin-, Cd31-, Sca+ cells. **(H)** Representative scatter plot of flow cytometry analysis of cells isolated from SM22-rtTA-cre;mTmG mice showing overlap between GFP+ cells (SM22+) with Lin-, Cd31-, Sca+, Pdgfrα+, Pdgfrβ+ cells. n=4-6, *p<0.05, **p<0.01, and ***p<0.001.

Single-cell analysis unveiled a shared origin for all adipogenic trajectories across both treatments, implicating mural cells expressing transgelin (SM22) (**Fig. 3C**). This cell subpopulation was distinguished from previously reported cell types. These mural cells lacked expression of Pdgfrα, while Pdgfrβ was expressed in numerous cell types (**Fig. S.3B**). Cd81, a recently reported marker, was expressed in all progenitor cell types/states expressing Sca1, including those expressing Pdgfrα and/or Pdgfrβ. Hence, to validate our findings in vivo, we used SM22-rtTA-cre;mTmG mural tracer mouse model. After seven days of doxycycline administration to induce membrane-bound GFP expression in SM22-expressing cells, followed by a three-day washout period, mice were either treated with CL− or exposed to cold for three days. Strikingly, only in the cold-exposed cohort was the co-staining of GFP with perilipin or Ucp1 evident (**Fig. 3D**), suggesting that while Adrb3 activation primes APCs, only cold triggers the differentiation of these primed APCs into beige adipocytes.

During the initial phase of adipogenesis, progenitor cells undergo mitotic clonal expansion before cell differentiation^29^. To assess the proliferation dynamics of SM22+ cells, we administered EdU (5-ethynyl-2′-deoxyuridine) to SM22-rtTA-cre;mTmG mice for 24 hours preceding their three-day regimen of either vehicle, CL−, or cold exposure. We then assessed Edu incorporation in Lin-, Cd31-, Sca+, and GFP+ cells using flow cytometry. Remarkably, a significant 85.8% of SM22+ cells incorporated EdU in CL− treated mice, compared to just 5.7% in vehicle-treated mice. In contrast, only 17.3% of GFP+ cells exhibited proliferation in response to cold exposure (**Fig. 3E**). The ratio of GFP+ APCs to total APCs (GFP+ + GFP-) remained consistent between the vehicle and CL− treated cohorts but decreased significantly in cold-exposed mice (**Fig. 3F**). This downward shift was not tied to a decline in APC numbers in cold-exposed mice since the total count of APCs increased similarly to CL− treated mice (**Fig. 3G**). Over a seven-day time frame, we recorded a gradual increase in proliferating SM22+ cells following CL treatment (**Fig. S3C**), indicating an ongoing expansion of this cell population, potentially due to the absence of a unique cold-dependent signaling pathway leading to the recruitment of proliferating cells.

Earlier studies have shown that all progenitor cells belong to the Pdgfrα+ lineage in subcutaneous adipose tissue. These cells can differentiate into two distinct preadipocyte types: Pdgfrβ−, which has the potential to differentiate into beige adipocytes, and Pdgfrβ+, which can differentiate into unilocular white adipocytes^26^. We hypothesized that SM22+ cells give rise to APCs. Eight-week-old SM22-rtTA-cre;mTmG mice were administered doxycycline for four weeks to label a substantial proportion of APCs. Flow cytometry analysis revealed both Pdgfrα+ and Pdgfrα+/Pdgfrβ+ cells exhibiting GFP expression (**Fig. 3H**), suggesting that the majority of APCs in WAT originate from SM22+ mural cells in adult mice.

Next, we aimed to investigate the mechanisms driving the differentiation of SM22+ mural cells into beige adipocytes under cold exposure. Given that both cold and CL− treatments prime APCs in vivo, we postulated that a differentiation-specific signal might be absent in the CL− treated group. To uncover this early signal, we performed RNA sequencing on isolated progenitor cells from mice treated with an Adrb3 agonist or a vehicle for 24 hours. Our analysis identified 345 upregulated genes while only four genes were downregulated (**Fig. 4A**). As expected, functional enrichment analyses revealed pathways associated with glucose and lipid metabolism alongside mitochondrial function-related pathways were significantly upregulated post-CL− treatment (**Fig. 4B**). Notably, Adrb1 emerged as one of the most prominently upregulated genes. While Adrb1 has previously been recognized for its role in brown and beige fat activation^49,50^, recent investigations found its contribution to the thermogenic and lipolytic functions of mature brown and beige adipocytes to be limited^51^. Interestingly, we observed an induction of Adrb1 expression in APCs harvested from mice treated with either CL− or cold exposure for three days (**Fig. 4C**). Despite the relatively low detectable levels of Adrb1 in single-cell data, which may be attributed to the limitations of sequencing depth, pathways linked to adrenergic receptor signaling and cAMP metabolism were distinctly enriched across several progenitor cell types following cold or CL− exposure (**Fig. 4D**).

**Figure 4:**
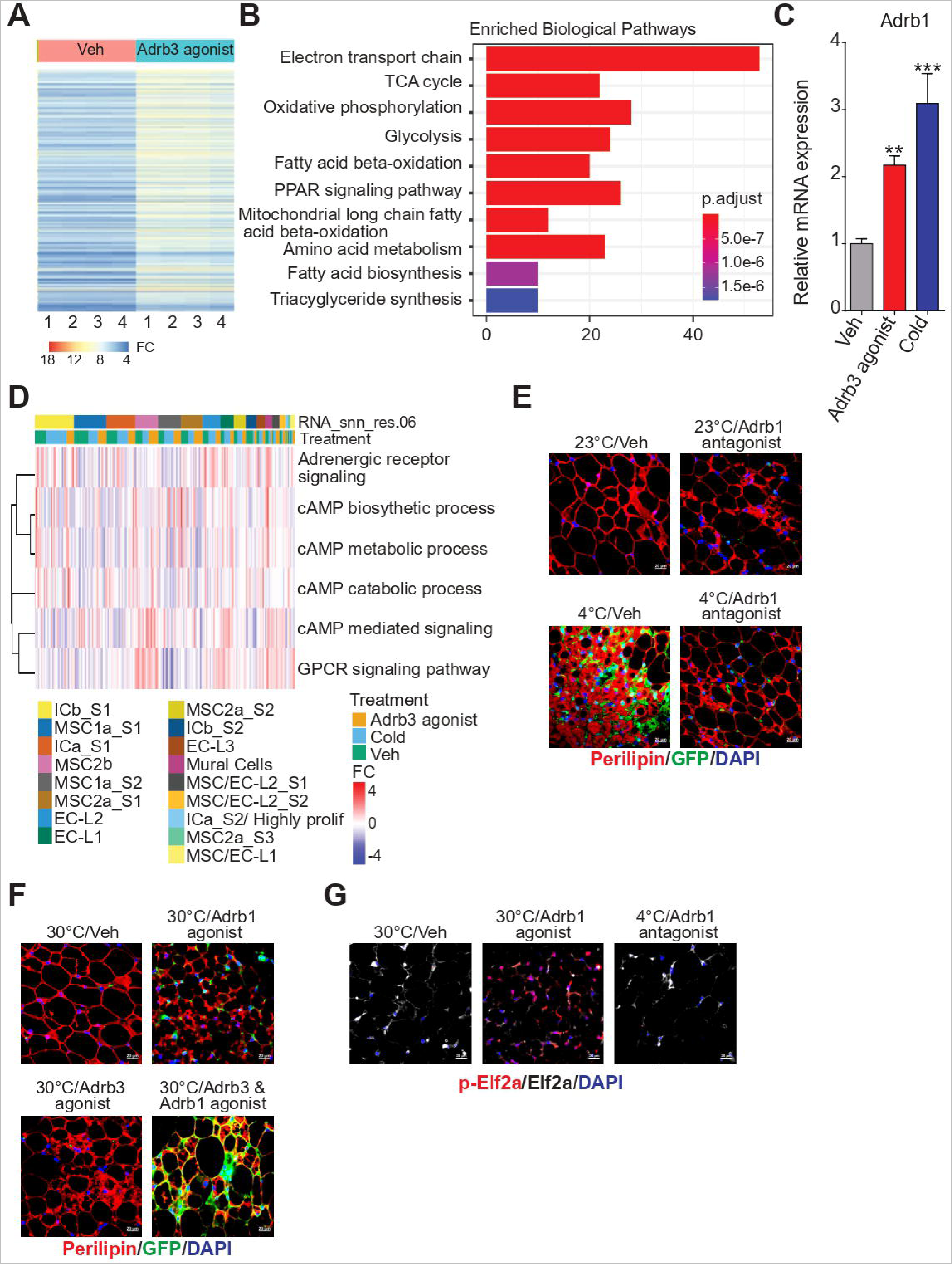
Beige Progenitor Cell Recruitment Requires Activation of Both Adrb1 and Adrb3. **(A)** FACS analysis of EdU incorporation in GFP+, Lin-, Cd31-, Sca+ cells following 7-days CL316,246-time course injections in vivo (mice received a single dose daily, n=5). **(B)** RNAseq heatmap of Lin-Sca+ cells isolated following 24h treatment with CL316,246 (1 mg/kg) **(C)** bar graph of enriched biological pathways in Lin-Sca+ cells isolated following 24h treatment with CL316,246. **(D)** Gene expression of Adrb1 in Lin-Sca+ cells. **(E)** Heatmap of Go terms associated with adrenergic receptor signaling. **(F)** Immunostaining in SM22-rtTA-cre;mTmG mice housed either housed at room temperature (RT) or at 4°C and treated with either saline (veh) or Adrb1 antagonist (betaxolol hydrochloride, 5 mg/kg) for 3 days. Perilipin (Red), GFP (green). **(G)** Immunostaining in SM22-rtTA-cre;mTmG mice housed either housed at room temperature (RT) or at 30 °C and treated with either treated saline (veh) or Adrb1 agonist (denopamine, 4 mg/kg), CL316,246 (1 mg/kg) or a combination of both for 3 days. Perilipin (Red), GFP (green). **(H)** Immunostaining of p-elf2a and ef2a in mice housed either at 30°C or 4°C and treated with either Adrb1 antagonist (betaxolol hydrochloride, 5 mg/kg) or Adrb1 agonist (denopamine, 4 mg/kg). n=4-6, *p < 0.05, **p < 0.01, and ***p < 0.001.

This observation prompted us to hypothesize that Adrb1 activation is crucial for the differentiation of SM22+ APCs into beige adipocytes. To test our hypothesis, SM22 lineage-tracing mice were subjected to cold exposure while being treated with a selective Adrb1 antagonist (betaxolol hydrochloride) for three days. No differences in shivering were observed between the control and treated groups. Histological analysis showed that the in vivo Adrb1 inhibition resulted in an elimination of SM22+ APC recruitment in cold-exposed mice, thereby mirroring the effect of CL− at room temperature (**Fig. 4E**). Furthermore, SM22 lineage tracing mice, housed at thermoneutrality for four weeks and co-treated with an Adrb3 agonist (CL−) and an Adrb1 agonist (denopamine), showed a synergistic effect that effectively mimicked the cell recruitment observed with cold exposure, an effect not observed with either treatment alone (**Fig. 4F, S4A**).

To complement our findings, we performed single-cell RNA sequencing on lineage-negative stromal cells from adult mice’s inguinal white adipose tissue (WAT), post-treatment with denopamine alone or combined with CL− for three days (**Fig. S4B**). Expectedly, neither treatment alone altered the cell population compared to the vehicle, cold, or CL− treatments (**Fig. S4C**). Interestingly, the combination of denopamine and CL treatments mirrored the cold-induced changes in cell population numbers (**Fig. S4D**). Moreover, the co-activation of Adrb1 and Adrb3 in SM22+ lineage tracing mice, housed at thermoneutrality, led to SM22+ cell recruitment into beige adipocytes, at levels comparable to cold exposure (**Figs. 4F, 3D**). Since compared to CL cold induces p-Elf2a we wondered whether activation of Adrb1 was sufficient to promote its phosphorylation. When housed in thermoneutrality, we found that Adrb1 activation resulted in a significant upsurge in p-Elf2a phosphorylation, while Adrb1 inhibition at 4C hindered its phosphorylation (**Fig. 4G**). Collectively, our data underscore the essential, yet distinct roles of Adrb1 and Adrb3 activation in the beige adipogenesis of SM22+ APCs. While Adrb3 activation is necessary to promote Adrb1 upregulation within progenitor cells, Adrb1 activation, in turn, is crucial for the activation of the unfolded protein response (UPR) and subsequent differentiation into mature beige adipocytes.

Consistent with previous studies we found that Adrb3 expression is restricted to mature white adipocytes and is not detected within the SVF^30^ (**Fig. 5A**). To elucidate the link between Adrb3 activation in mature adipocytes, APCs priming, and cell proliferation, we performed an RNA-seq on mature adipocytes isolated from mice receiving a single 24-hour CL− treatment. We found that 71 genes were significantly upregulated by CL treatment while 269 genes were downregulated (**Fig. 5B**). As expected, pathways enrichment analysis of upregulated genes showed an association with fatty acid metabolism, tricarboxylic acid (TCA) cycle, respiratory electron transport, and triglyceride metabolism. Interestingly, among the top enriched pathways were terms such as α−linolenic (omega3) and linoleic (omega6) acid metabolism and α−linolenic acid (ALA) metabolism. Linoleic acid (LA, 18:2, n-6) and α-linolenic acid (ALA, 18:3, n-3) are the most abundant 18 carbon (C18) essential long-chain polyunsaturated fatty acids (PUFA). Prior studies have proposed that diets rich in n-3 PUFA reduce obesity, diminish fat deposition, decrease fasting serum triglyceride concentrations, and boost energy expenditure and fatty acids oxidation in human and rodent studies^31,32^. Recent research has suggested that PUFA from dietary fat is the lipolytic mediator for these effects in brown and white fat depots^33^. These data indicate that lipolysis-derived PUFAs could be instrumental in the early reprogramming of APCs via Adrb3-activated lipolysis in mature adipocytes. To test this hypothesis, we employed Acipimox, an FDA-approved hormone-sensitive lipase (HSL) inhibitor, to assess whether lipolysis influences SM22+ APCs recruitment.

**Figure 5:**
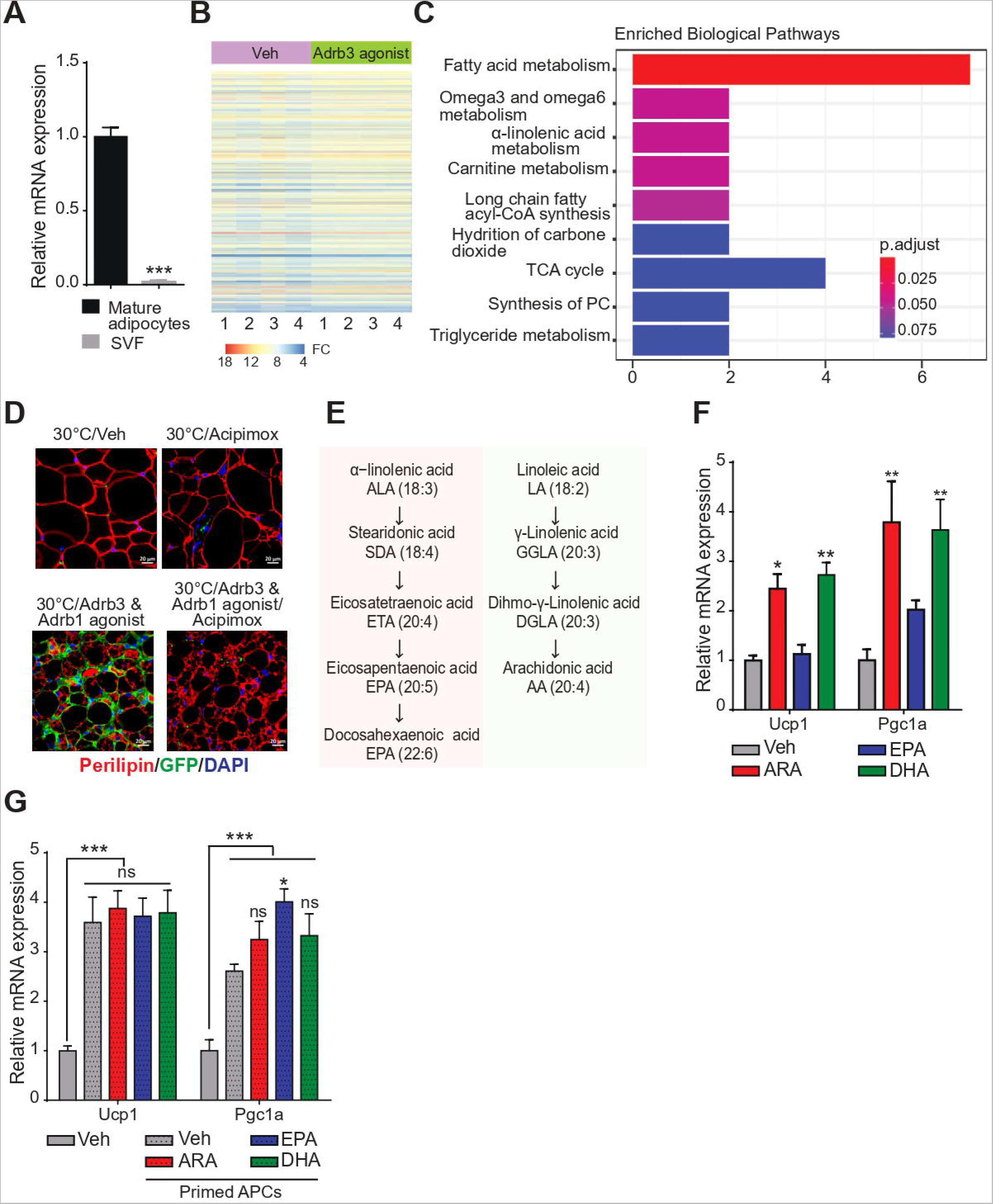
Lipolysis-Derived Eicosanoids Promote Beige Priming in SM22-Expressing Mural Cells. **(A)** Gene expression of Adrb3 in mature adipocytes and stromal cells from inguinal depot. **(B)** RNAseq heatmap of Adipocytes isolated following 24h treatment with CL316,246 (1 mg/kg). **(C)** bar graph of enriched biological pathways in Adipocytes isolated following 24h treatment with CL316,246. **(D)** Immunostaining in SM22-rtTA-cre;mTmG mice housed at thermoneutrality (30°C) for 4 weeks and treated with single dose of lipolysis inhibitor Acipimox (50 mg/kg) alone or in combination with Adrb1 agonist (denopamine, 4 mg/kg) and Adrb3 agonist (CL316,246 1 mg/kg) for 3 days. **(E)** Schematic of the lipolytic metabolism of α-Linolenic Acid and Linoleic Acid **(F, G)** Gene expression from differentiated APCs isolated from mice treated with saline (F) or CL316,234 (G) for 24 h and pretreated with BSA conjugated arachidonic acid (ARA, 10 nM) or eicosapentaenoic (EPA, 10 nM) or docosahexaenoic acid (DHA, 10 nM) for 2 days. n=4-6, *p < 0.05, **p < 0.01, and ***p < 0.001.

Considering the lethality of lipolysis inhibition under cold exposure, we used SM22 lineage tracing mice housed at thermoneutral conditions. These mice were subjected to a combination of Adrb1 and Adrb3 agonists to promote SM22+ APCs recruitment, either with or without an anti-lipolytic agent for three days. We found that the recruitment of SM22+ APCs into beige adipocytes was abolished by Acipimox treatment (**Fig. 5D, S5A**), implying a vital role for free fatty acid (FFA) release in the beige adipogenesis of vascular progenitors.

Metabolism of ALA and LA generates multiple bioactive intermediaries including eicosapentaenoic acid (EPA), docosahexaenoic acid (DHA), and arachidonic acid (AA) respectively (**Fig. 5E**). To further decipher the signaling role of FFA, we treated APCs with BSA-conjugated EPA, DHA, and AA. Remarkably, AA and DHA treatment induced APC priming into beige adipocytes, while EPA showed less potency (**Fig. 5E**). This effect was not observed when APCs from mice primed with β3-AR were treated with different PUFA species (**Fig. 5F**), suggesting that the products of LA and ALA lipolysis are the mediators Adrb3 induced SM22+ APCs reprograming and recruitments to beige adipocytes.

## DISCUSSION

While it was known that young adult WAT harbors precursor cells with the capacity to undergo beige adipocyte differentiation^20,21,24,25,26,27,23^ our study identified a new mechanism governing the emergence of beige adipocytes during cold exposure. We also revealed a profound reprogramming of pre-existing adipogenic trajectories towards thermogenic phenotypes, rather than reliance on a distinct beige precursor cell. We found that, irrespective of the prompting factor for WAT beiging, the progenitor cell populations were unaltered. Intriguingly, we observed a notable redistribution of cell types contributing to diverse WAT lineages. Our investigation revealed that a significant proportion of these cells could be traced to a distinct naïve mural cell population expressing SM22.

While previous research established that committed preadipocytes originate from the Pdgfrα+ lineage and can shift from a beige to a white progenitor state driven by Pdgfrβ+ expression^26^, our findings indicate that both Pdgfrα+ and Pdgfrα+/Pdgfrβ+ progenitor cells are derived from the SM22+ lineage. Despite earlier suggestions of a beige-specific progenitor cell^23,34^, such as Cd81-expressing cells, we found that Cd81 is broadly expressed in Lin-stromal cells and significantly overlapping with Sca+ cells, including Pdgfrα+ and/or Pdgfrβ+ cells, pointing towards their derivation from the SM22+ lineage. Similarly, although Bts2^high^ cells have been proposed as a unique beige progenitor, our research indicates its expression across multiple cell types, including endothelial cells (EC2), mesenchymal cells (MSC2b), and mural cells, some of which originate from SM22+ cells (**Table 1**). It is possible that cells become differentially permissive to beige reprogramming or priming as they progress along specific lineages and transitionally enriching for cells expressing markers such as Cd81 or Bts2.

The sequential downregulation of Pdgfrα and Pdgfrβ determines the thermogenic phenotype upon differentiation, highlighting their importance^26,27,35^. While further investigations are warranted to discern the extent of cells’ permissibility to beige reprogramming and whether this process is altered with obesity or aging, our findings reveal in vivo priming of progenitor cells in response to cold exposure and Adrb3 agonist treatment. Furthermore, the preservation of beige priming during isolation, expansion, and in vitro differentiation indicates that progenitor cells integrate thermogenic signals to activate the beige phenotype as they undergo the adipogenic commitment. Clarifying the differences between physiological and pharmacologically induced beiging mechanisms provides essential insights, particularly in the context of physiological Adrb3 activation versus the induction via Adrb3 agonists. Recent evidence suggests that beige adipocytes emerge through transdifferentiation of white mature adipocytes upon Adrb3 stimulation, while cold exposure facilitates both beige transdifferentiation and progenitor cell recruitment^36,20^. Our study highlights that both treatments promote progenitor cell priming and expansion. However, a relatively restrained proliferation of marked SM22+ cells from mice exposed to cold, indicates that a fraction of these cells underwent beige adipogenesis. Consistent with prior findings, our histological analysis demonstrated that only cold exposure triggers the recruitment of marked SM22+ cells into mature beige adipocytes.

The progression of progenitor cells into mature beige adipocytes is orchestrated by an array of signaling and transcriptional programs that facilitate sequential fate determination and the acquisition of the adipogenic phenotype^37^. Our investigation identified five pivotal pathways underpinning these programs. We ascertained that beige phenotype progression involves the downregulation of genes related to extracellular matrix organization and antigen processing & presentation^10,38,39,40,41,42,43^. In addition, our data highlight the upregulation of fatty acid transport, actin cytoskeleton reorganization, and response to unfolded protein (UPR) as characteristic hallmarks of progenitor cell progression toward the beige lineage. Prior studies have drawn associations between genes within these pathways and the thermogenic capacity of adipose tissue suggesting the potential for manipulating these pathways to enhance beige adipogenesis^44,45,46,47^. A particularly intriguing observation pertains to the activation of UPR in response to cold exposure. This aligns with recent research proposing the essential role of PERK/eIf2a activation in mitochondrial thermogenesis within BAT^48^.

Wherein both Adrb3 agonist and cold exposure lead to progenitor cell priming and proliferation, while only cold exposure triggers beige adipocyte adipogenesis, strongly implied the involvement of distinct signaling cascades upon cold exposure that enables adipogenic progression. Using multiple approaches including transcriptomic analyses, pharmacological interventions, and in vivo fate-mapping techniques, we unveiled a pivotal role for sequential Adrb1 activation within progenitor cells. This tightly regulated activation sequence proves indispensable for primed SM22+ progenitors to exit the cell cycle and subsequently drive their differentiation into mature beige adipocytes. These findings align with prior reports emphasizing the substantial contribution of Adrb1 in promoting progenitor cell recruitment^36^.

Previous observations have suggested that Adrb1 could be important to promote progenitor cell proliferation^49,50^, however, a recent genetic study found that Adrb1 knockdown in progenitor cells does not influence cold-induced progenitor cells proliferation^51^. Our data demonstrate that Adrb1 activation is important for the cells to exit from the cell cycle and differentiate into beige adipocytes potentially mirroring the effect of the chemical agent IBMX, traditionally employed for in vitro differentiation of cell lines and primary cells into adipocytes. In line with reported findings in other cell types, such as cardiomyocytes, we observed Adrb1-mediated activation of UPR. This further supports the notion that Adrb1 activation promotes mature adipocyte differentiation and/or mitochondrial biogenesis through the activation of the UPR^52,53,54,48^. Further in-depth investigations are needed to better understand the interplay between those.

Our findings establish a previously unappreciated link between the activation of Adrb3 in mature adipocytes and the initial priming of SM22+ mural cells. Our investigation underscores the pivotal role of lipolysis products as central mediators orchestrating beige reprogramming within this context. Importantly, our data highlight that it is not the activation of Adrb3 per se that is important, but rather the bioactive molecules generated through lipolysis. It is pertinent to acknowledge that previous studies have reported that Adrb3 null mice still possess a potential for cold-induced beiging. However, the interplay of multiple non-adrenergic pathways inducing lipolysis, combined with potential mouse strain disparities, may account for the divergent outcomes observed^55,56,57,58^. A particularly distinctive aspect is that the experiments involving Adrb3 null mice were conducted at room temperature, while our investigations were undertaken at thermoneutrality using a combination of lipolysis inhibitor and Adrb1 & 3 agonists which enabled us to halt lipolysis completely.

Collectively, our work pioneers a novel model elucidating mechanisms governing cold-induced beige adipogenesis. Cold exposure triggers a cascade where lipolysis-derived products play a pivotal role in priming naïve mural SM22+ cells, subsequently driving their expansion. As these cells progress, they undergo a developmental trajectory, transitioning into PDGFRα+ and/or PDGFRα+/PDGFRβ+ mesenchymal cells boasting robust adipogenic potential. A defining feature of the beige lineage progression is the emergence of Adrb1 expression. The activation of Adrb1 assumes primordial importance, orchestrating the recruitment and differentiation of these cells into fully mature beige adipocytes.

These collective insights reveal a pivotal communication axis, underscoring the dynamic interplay between mature adipocytes and mural cells during the course of cold acclimation. Furthermore, the existence of a unique reservoir of naïve cells responsive to metabolic cues, and capable of diversifying into multiple lineages, emerges as a significant finding. This prompts the need for future investigations, particularly in the contexts of obesity and aging, to unveil the attributes of these cells and decipher how they seamlessly integrate signals emanating from the diverse array of cell types residing within the adipose tissue.

## ACKNOWLEDGMENTS

We thank Hu Tianmu and Yuriy Alekseyev of the Boston University School of Medicine (BUSM) Single Cell Sequencing Core for their advice and assistance. We thank Jessica E. C. Jones for the help with the RNA-seq data generation. We also thank Anna C Belkina and the BUMC Flow Cytometry Core Facility for their support and Eleni Anastasakou for the help with the graphic abstract. This project was funded by grant# U24DK097771 from the National Institute of Diabetes, Digestive, and kidney diseases via the NIDDK Information Network’s (dkNET) New Investigator Pilot Program in Bioinformatics (N. Rabhi), the National Institutes of Health/NIDDK (DK117161 and DK117163 to SR. Farmer,DK134534 to MD. Layne and SR. Farmer, and DK132080 to MD Layne), the Department of Biochemistry & Cell Biology at Early Career Award (N. Rabhi), The Genome Science Institute award (N. Rabhi, SR. Farmer).

## AUTHOR CONTRIBUTIONS

K Desevin: investigation, data collection.

BN Cortez: investigation, data collection.

JZ Lin: investigation, data collection.

D Lama: data collection.

MD Layne: resource and funding acquisition, review and editing

SR Farmer: conceptualization, funding acquisition, and writing—review and editing.

N Rabhi: conceptualization, data curation, bioinformatic data analysis, funding acquisition, and writing—original draft, review, and editing.

---

**Supplementary figure 1: Sustained Cellular Heterogeneity in WAT Unaltered by β3-Adrenergic or Cold Stimulation. (A)** Violin plots showing the expression levels and distribution of marker genes from the literature. (**B)** Individual genes t-SNE plot showing the expression levels and distribution of representative marker genes of each cluster. **(C)** t-SNE plot of iWAT Lin-stromal vascular cell types comparison from mice treated with vehicle, CL−316,243 or exposed to 4°C for 3 days.

**Supplementary figure 2: Shift of Cell Trajectory in Response to Cold Exposure and β3-Adrenergic Stimulation (A)** Violin plots showing the expression levels and distribution of representative marker genes of each cell type along different cell states. **(B)** Violin plots showing the expression levels and distribution of marker genes from the literature in each cell state. **(C)** Heatmap of the top differentially expressed genes. **(D)** Side-by-side t-SNE plots of iWAT progenitor cell types state from Vehicle, CL−316,243 and exposed to 4°C for 3 days. **(E)** t-SNE plot of progenitor cell types state comparison from Vehicle, CL−316,243 and exposed to 4°C for 3 days. **(F)** Bar plot of the average weight for each lineage in each condition. **(G)** Heatmaps of differentially expressed genes between conditions ordered according to hierarchical clustering on the cold exposed condition. **(H)** t-SNE plot showing scores of top gene ontology pathways (biological process) associated with genes differentially expressed in cold exposed samples.

**Supplementary figure 3: SM22 expressing mural cells are primed into the Beige adipogenic cells but only cold promote their Recruitment. (A)** Schematic illustration of the experimental design. **(B)** t-SNE plot of expression levels and distribution of indicated genes. **(C)** Quantification of flow cytometry data analysis of Edu incorporation in GFP+ Lin-, Cd31-, Sca+ cells following 7-days CL316,236-time course daily injection in vivo (n=5). *p < 0.05, **p < 0.01, and ***p < 0.001.

**Supplementary figure 4: Beige Progenitor Cell Recruitment Requires Activation of Both Adrb1 and Adrb3. (A)** Immunostaining in SM22-rtTA-cre;mTmG mice housed either housed at room temperature (RT) or at 4°C and treated with either saline (veh), Adrb1 antagonist (betaxolol hydrochloride, 5 mg/kg) or Adrb1 agonist (denopamine, 4 mg/kg) for 3 days. **(B)** t-SNE plots showing different Lin-stromal vascular cell types in iWAT isolated from mice treated with Vehicle, cold, betaxolol hydrochloride (Adrb1 agonist), CL−316,243 (Adrb3 agonist) or both adrb1 and Adrb3 agonists for 3 days. (C) Side-by-side t-SNE plots of iWAT progenitor cell types state from Vehicle, CL−316,243 and exposed to 4°C for 3 days. (D) Bar plot showing the percentage of cells per cluster in each treatment.

**Supplementary figure 5: Lipolysis-Derived Eicosanoids Promote Beige Priming in SM22-Expressing Mural Cells: (A)** Immunostaining in SM22-rtTA-cre;mTmG mice housed at thermoneutrality (30°C) for 4 weeks and treated with single dose of lipolysis inhibitor Acipimox (50 mg/kg) alone or in combination with Adrb1 agonist (denopamine, 4 mg/kg) and Adrb3 agonist (CL316,246 1 mg/kg) for 3 days.

## METHOD DETAILS

### Animals

All animal studies were approved by the Boston University Chobanian and Avedisian School of Medicine Institutional Animal Care and Use Committee. C57BL/6 mice were purchased from The Jackson Laboratory (JAX) at 6 weeks of age and acclimated for 2 weeks. TetO-Cre (JAX #006234) mice were bred with ROSAmT/mG (JAX #007676) to generate TetO-Cre; ROSAmT/mG mice. These mice were crossed with SM22-rtTA (JAX #006875) to generate SM22-rtTA-cre; TetO-Cre;mTmG. Mice were housed in a temperature-controlled environment with a 12 hr light-dark cycle and ad libitum water and a standard chow diet. When specified, mice were housed in thermoneutral chambers at 30°C. To induce Cre expression, male mice were fed a doxycycline diet (TD.120769, 625 mg/kg) for 7 days followed by a 3 days washout before any treatment. For the treatment with vehicle (saline), CL−316,243 (Sigma-Aldrich, 1 mg/kg), Denopamine (Sigma-Aldrich, 4 mg/kg), Betaxolol hydrochloride (Sigma-Aldrich, 5 mg/kg) or Acipimox (Sigma-Aldrich, 50 mg/kg) mice were injected intraperitoneally (i.p.) for 3 consecutive days. For the cold exposure experiment, mice were maintained in a 4°C room for 3 days.

For 5-ethynyl-2‘-deoxyuridine (EdU) labeling of proliferating cells, mice were injected with EdU (Invitrogen, 20 mg/kg, i.p.) prepared in sterile PBS. EdU was administered once 24 hours before euthanasia. After isolation, cells were analyzed as indicated by the manufacturer.

### Cell Isolation

Inguinal white adipose tissues (iWAT) from control and treated mice were collected and processed for isolation of stromal vascular cells (SVC) using the mouse adipose tissue dissociation kit (Miltenyi Biotec) and the gentleMACS™ Dissociator (Miltenyi Biotec) according to manufacturers instructions. When indicated in the figure legend MACS Anti-CD45 MicroBeads, for mice (130-052-301, Miltenyi Biotec) and MACS LD columns (Miltenyi Biotec) were used to deplete lineage+ (Lin+) cells and Anti-Sca-1 (non-HSC) MicroBeads (130-106-641, Miltenyi Biotec) for mice for enriching with Sca+ cells. For cell culture, cells were seeded with primary cell culture medium (DMEM/Ham’s F-12 with 10% FBS and 1% pen strep). The medium was replenished every 2–3 d. For adipogenesis, cells were incubated in an induction medium containing 1 μM dexamethasone, 870 nM insulin, 0.5 mM IBMX, and 1 μM rosiglitazone for 2 d. Cells were maintained in a differentiation medium containing the same concentrations of insulin afterward.

### Flow cytometry analysis and data processing

Freshly isolated iWAT SVC were resuspended in FACS buffer (PBS/1% BSA). Samples were blocked with mouse Fc block (BD Biosciences; 1:50) for 5 min then incubated with antibody mix supplemented brilliant stain buffer (BD Biosciences) and monocyte blocker (Biolegend) for 20 min at 4°C protected from light. SVC suspension was rinsed 3 times before flow cytometry analysis with Aurora spectral cytometry analyzer (Cytek Biosciences). The following antibodies were used Anti-mouse-CD45.2-BUV737(1:40, 564880, BD Biosciences), Anti-CD31-PECAM-1-eFluor450 (1:500, 48-0311-82, eBioscience), Anti-CD34-eFluor 660 (1:50, 50-0341-82, eBioscience), Anti-Sca-Ly6A/E-BUV395 (1:150, 563990, BD Biosciences) Anti-CD140a-Pdgfra-BV421 (1:300, 566293, BD Biosciences), and Anti-CD140b-Pdgfrb-PE-Cynine7 (1:250, 25-1402-80, Invitrogen). All data analyses were performed using FlowJo software (version 10.8.1). Data were plotted and compared using Prism 6.0 (GraphPad).

### Immunofluorescence

Paraffin-embedded iWAT were sections (5 µm) were mounted onto slides, deparaffinized, and rehydrated before performing antigen retrieval. Tissue sections were stained with rabbit anti-UCP1 (ab209483; Abcam, 1:500), rabbit perilipin-1 (#9349; Cell Signaling, 1:1000), rabbit anti-eIF2α (D7D3) (#5324, Cell Signaling), rabbit phospho-eIF2α (Ser51) (D9G8) (#3398, Cell Signaling), and, chicken anti-GFP (#1020, Aveslab) overnight at 4C. After washing with 0.1% tween-20 TBS, sections were incubated for 1 hr at room temperature with fluorophore-conjugated secondary antibody (donkey anti-chicken Alexa Fluor 488, donkey anti-rabbit Alexa 647, donkey anti-rabbit Cy3 (Invitrogen). Slides were then washed three times with 0.1% tween-20 TBS at room temperature in the dark. Coverslips were mounted using Prolong gold antifade (Thermofisher). Fluorescent images for all stained adipose tissue sections were captured with an Axio scan Z1 imager (Zeiss) at 63x magnification.

### Real-Time PCR

Total RNA was extracted from frozen tissues and cells using TRIzol reagent according to the manufacturer’s instructions. RNA concentrations were determined on NanaDrop spectrophotometer. Total RNA (100 ng to 1 µg) was transcribed to cDNA using Maxima cDNA synthesis (Thermo Fisher Scientific). Quantitative real-time PCR was performed on the ABI VIA detection system, and relative mRNA levels were calculated using the comparative threshold cycle (CT) method. SYBR green primers are listed in Table S2.

### RNA-seq

cDNA libraries for RNA-seq analysis were prepared from total RNA (isolated as described above) according to Illumina TruSeq stranded mRNA sample prep LS protocol. Pooled sample libraries were sequenced on Nextseq. 500 instruments with Nextseq. 500/550 High output kit, 150 cycles (Illumina).For data analysis, raw sequencing reads were subjected to quality control analysis via FastQC, and reports were aggregated via MultiQC^59^. Reads were then trimmed based on a sliding window and the presence of adapter sequences using Trimmomatic set to the following parameters: ILLUMINACLIP::2:30:10 LEADING:3 TRAILING:3 SLIDINGWINDOW:4:15 MINLEN:50^60^. Salmon was used to quantify the expression of transcripts to a reference transcriptome containing protein-coding and long non-coding RNAs from the gencode M10 annotation before aggregation to the gene level^61^. Only genes with a raw count of greater than 5 and present in at least half of the replicates were used for downstream analysis. Normalization and differential expression were performed using DESeq2 set to default parameters^62^. The biological replicates for each condition were pooled for differential expression analysis. Genes with a q-value of < 0.05 were considered significantly differentially expressed. GSEA pathway enrichment analysis was conducted on differentially expressed gene lists that were ranked by logFCLists of differentially expressed genes^63^.

### Single Cell RNA Sequencing

Cells were prepared for single-cell sequencing according to the 10x Genomics protocols. Sequencing was performed on Illumina NextSeq500. The Cell Ranger Single-Cell Software Suite (v.3.1.0) (available at https://support.10xgenomics.com/single-cell-gene-expression/software/pipelines/latest/what-is-cell-ranger) was used to perform sample demultiplexing, barcode processing, single-cell 3′ counting, and counts alignment to mm10 mouse reference genome. For further analysis, the R (v.3.1) package Seurat was used (adapted workflow available at https://satijalab.org/seurat/v3.1/immune_alignment.html)^64,65^. Briefly, cells with a feature count over 2500 or less than 200 or have over 5% of mitochondrial genes were filtered out. All the samples were integrated, and the top 18 dimensions were used to generate the final clusters. To distinguish between stem and progenitor cells the difference between the G2M and S phase scores was regressed out. Cells are represented with t-distributed stochastic neighbor embedding (t-SNE) plots. The Seurat function “FindNeighbors” followed by the function “FindClusters” were used for clustering. FindAllMarkers function was used to identify specific gene markers for each cluster. Violin plots were used to compare selected gene expressions. Differential expression between clusters was obtained using MAST. Specific genes for each cluster were used for functional annotation and Go terms using fgsea and pseudotime were analyzed using Slingshot^66^.

### Statistical analysis

Data were analyzed using GraphPad Prism software (GraphPad) or R software and are presented as mean +/- standard error of the mean (SEM). Statistical significance was determined by unpaired two-tailed Student t-test or ANOVA; a p-value of ≤ 0.05 was considered significant.

### Code availability

Codes are publicly available in the relevant citations and custom script is available on request.

