## Supplementary figures and images for "Adrenergic Reprogramming of Preexisting Adipogenic Trajectories Steer Naïve Mural Cells Toward Beige Differentiation"

### Supplemental figure 1

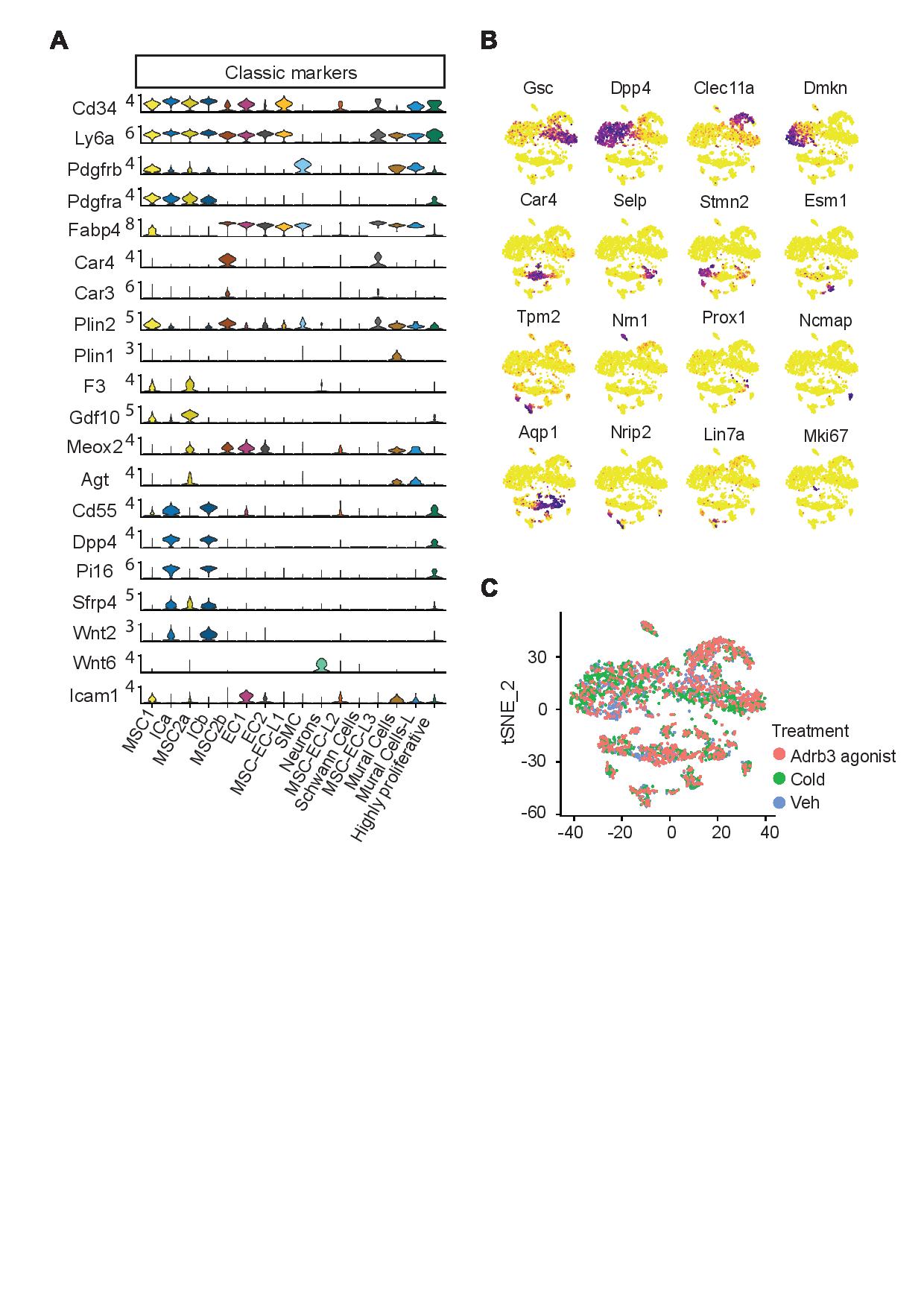

### Supplemental figure 2a

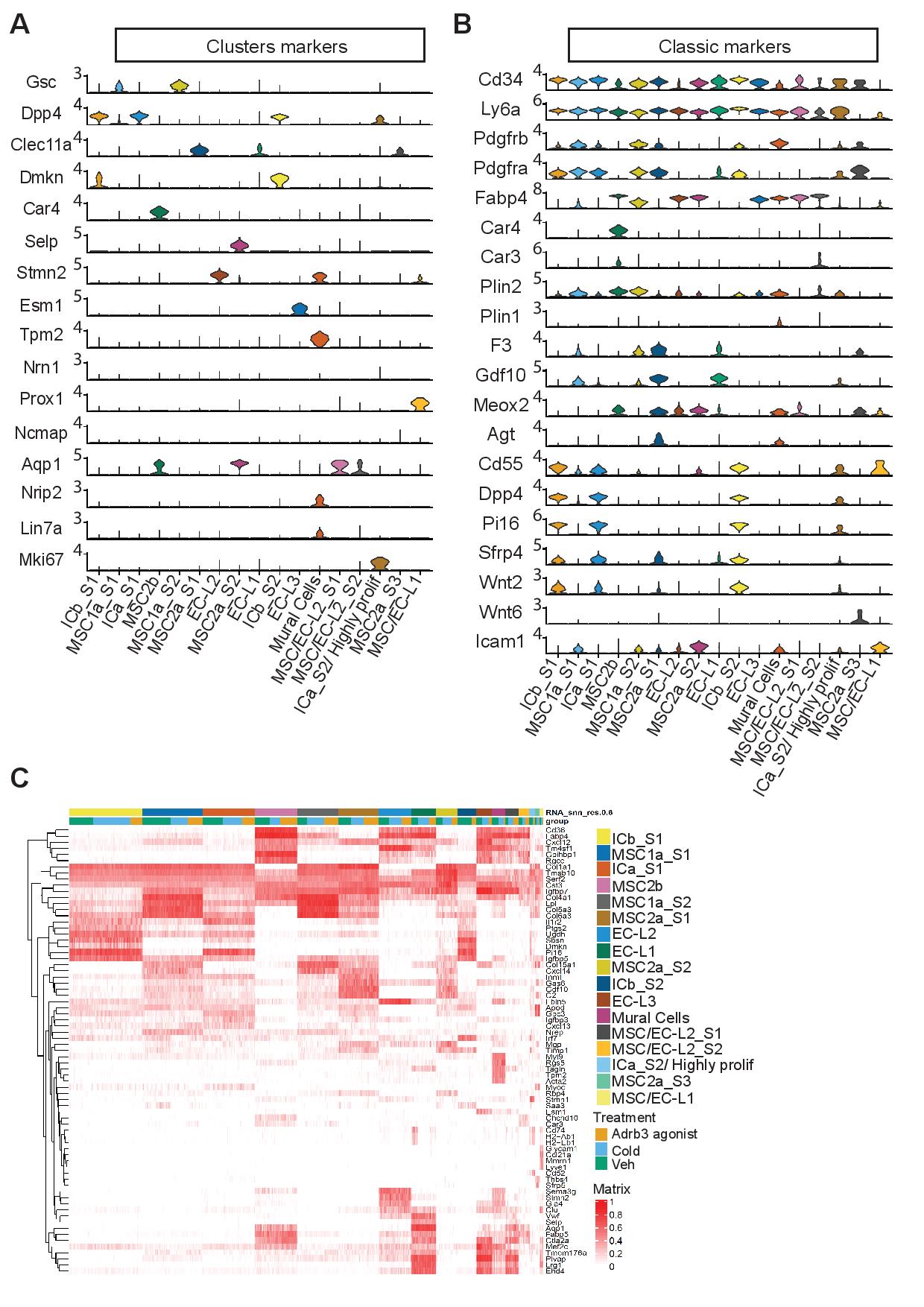

### Supplemental figure 2b

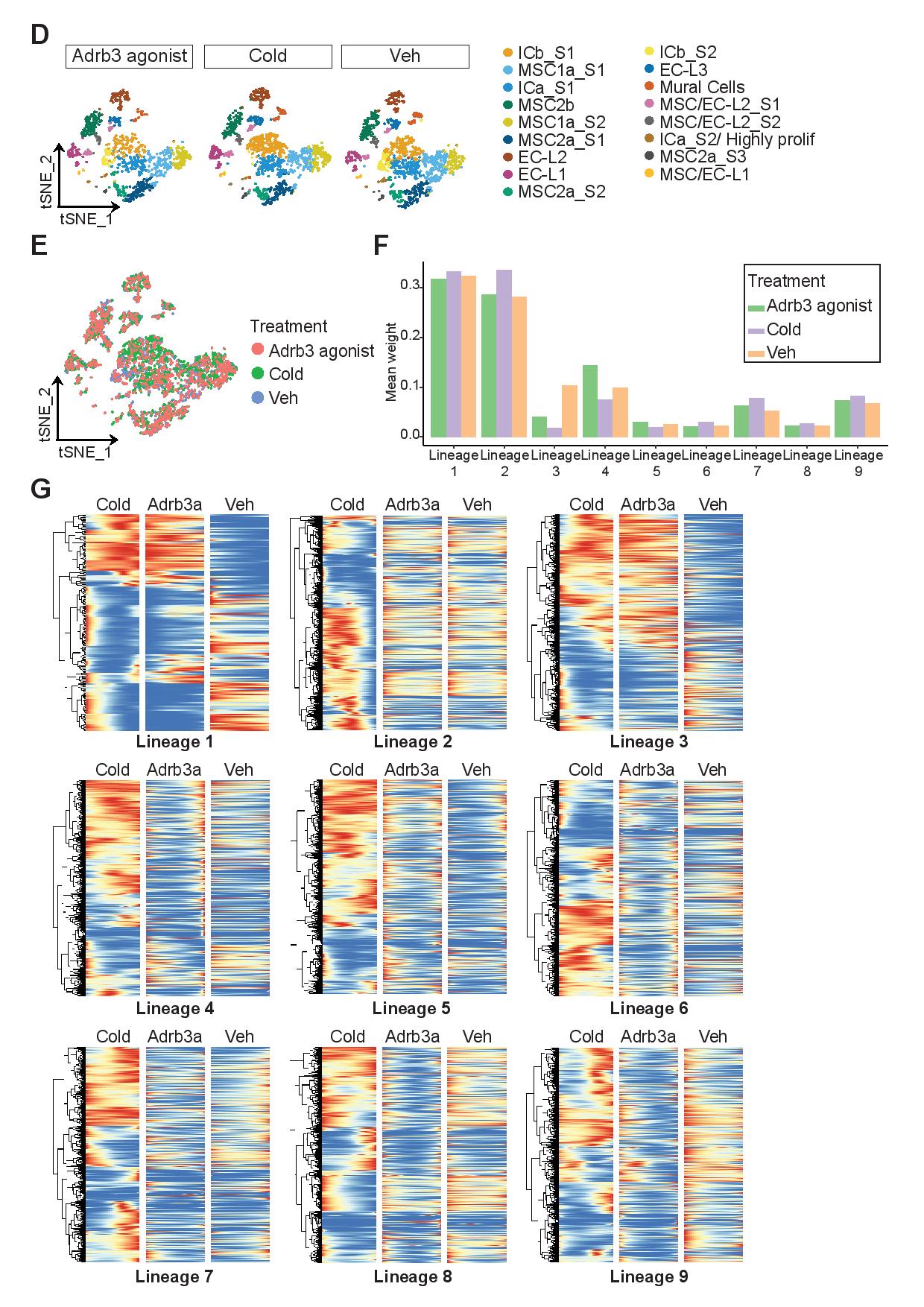

### Supplemental figure 2c

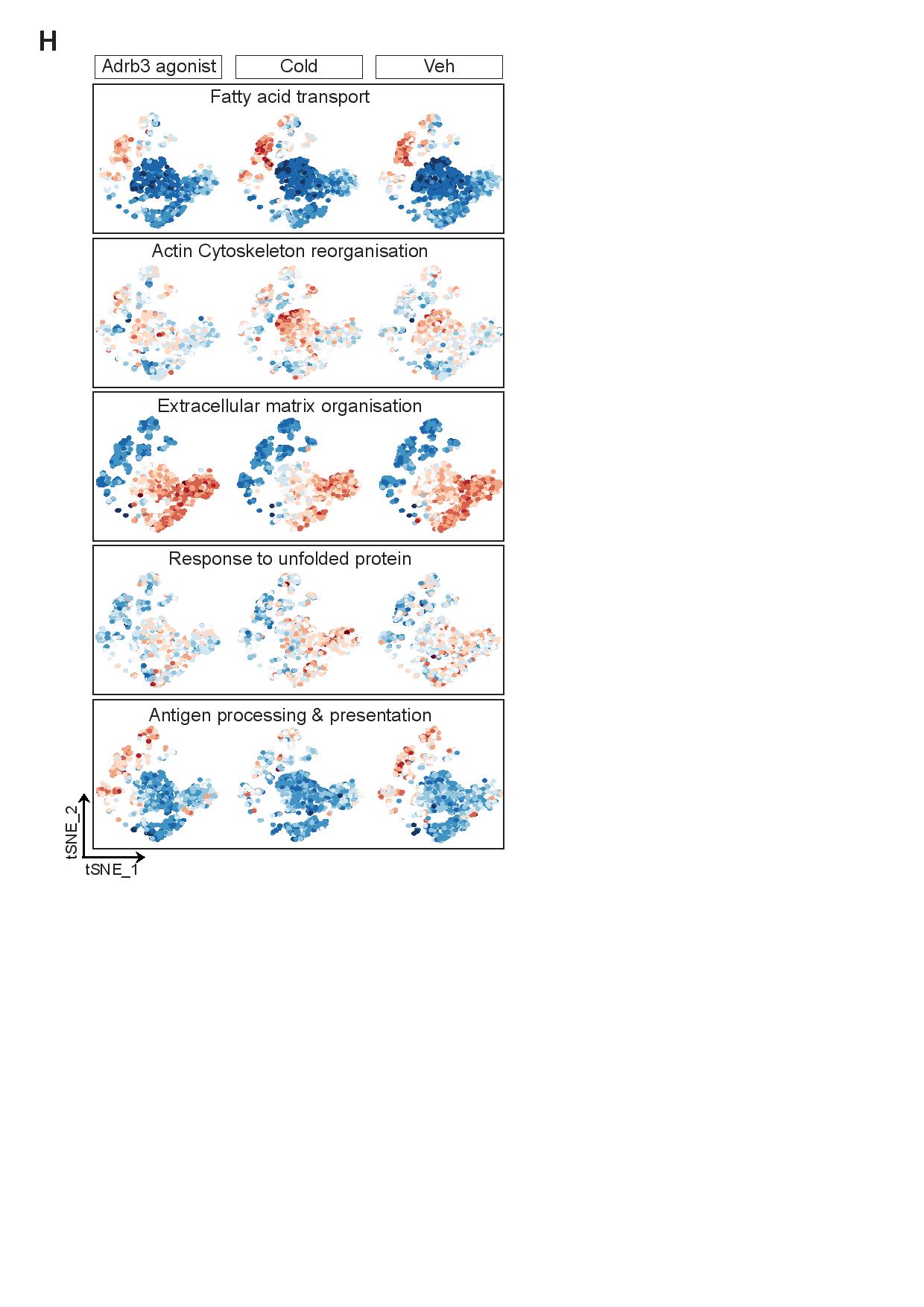

### Supplemental figure 3

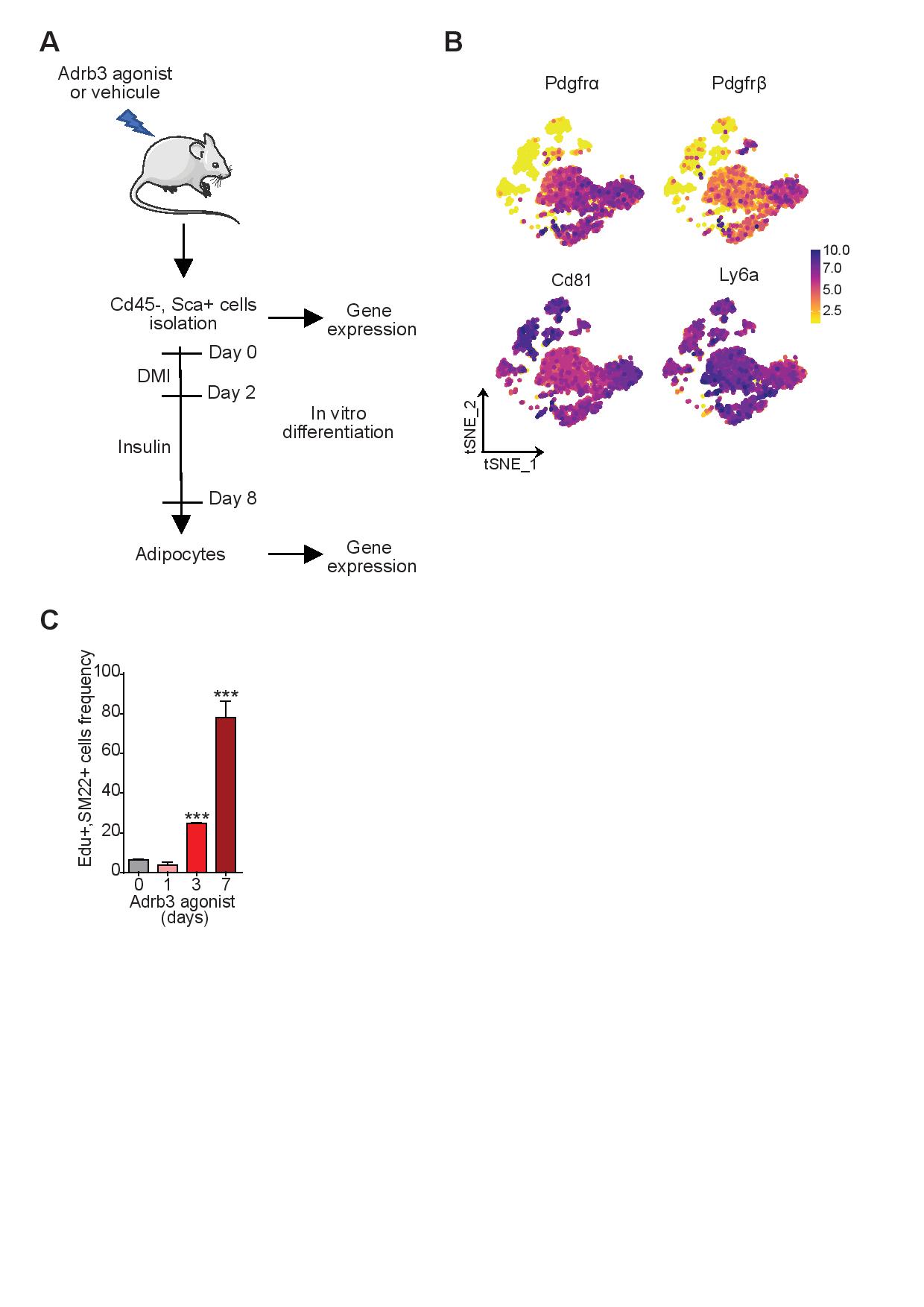

### Supplemental figure 4

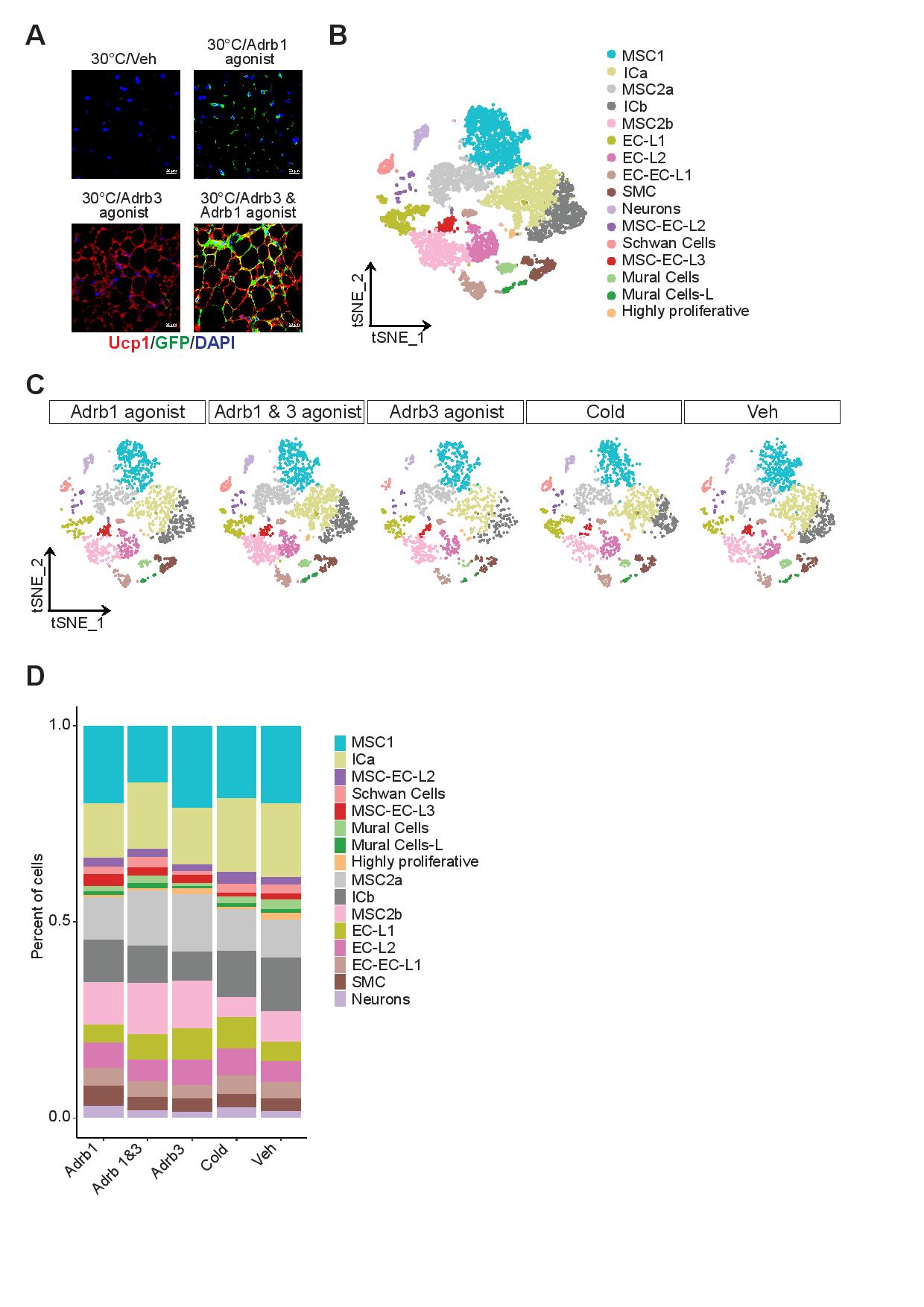

### Supplemental figure 5

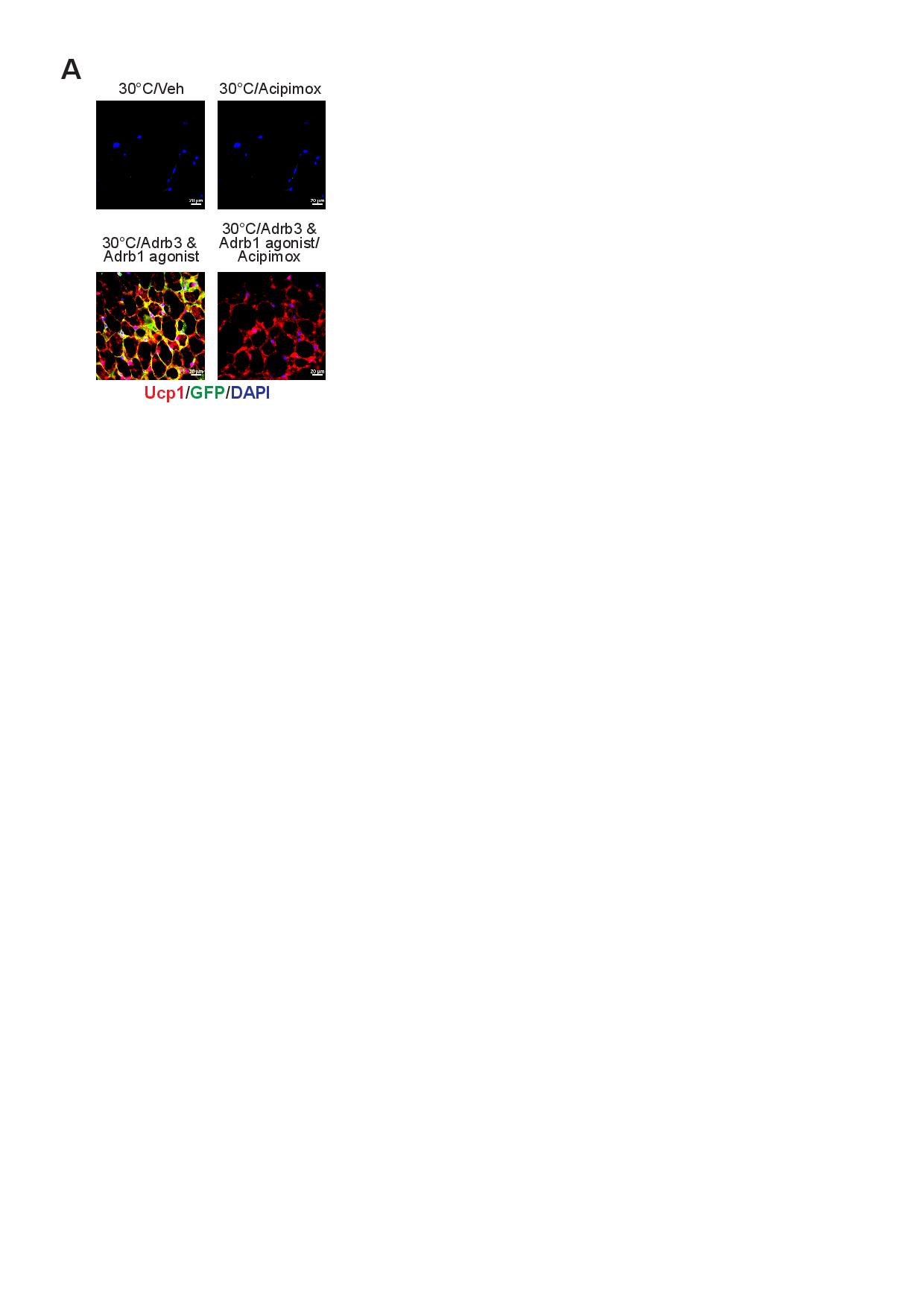
